# Discovery of populations endemic to a marine biogeographical transition zone

**DOI:** 10.1101/2020.07.25.221200

**Authors:** Tirupathi Rao Golla, Leishe Pieterse, Candice M. Jooste, Peter R. Teske

## Abstract

**Aim:** Biogeographical transition zones are areas of overlap between the faunas of adjacent biogeographical entities. Particularly, the well-defined transition zones along linear coastlines are interesting natural laboratories to study dispersal and incipient speciation. Few studies have explored whether marine biogeographical transition zones harbour biodiversity that is distinct from that of the biogeographical entities they separate. The Wild Coast in eastern South Africa is a poorly-studied transition zone between region’s warm-temperate and subtropical faunas, and is generally considered to be an area of faunal overlap.

**Location:** The South African portion of the Western Indian Ocean

**Methods:** Sequences of the DNA barcoding marker COI were generated from 306 estuarine sandprawns (*Callichirus kraussi*) collected at 13 sites. Genetic structure and evolutionary history were assessed using a haplotype network and a Bayesian discrete phylogeographic analysis.

**Result:** Two populations were identified whose ranges are centred on the Wild Coast, a rare one in the northern portion and a more common one in the central and southern portion of this biogeographical transition zone. These populations are not closely related to each other, but descend from subtropical and warm-temperate sister populations, respectively.

**Conclusions:** This is the first study to indicate that the Wild Coast marine biogeographical transition zone is not merely an area of faunal overlap, and one of very few studies to have discovered genetically unique populations within a marine biogeographical transition zone. The Wild Coast may harbour additional unique biodiversity that remains to be discovered, including rare species that require protection. More research is required to understand how this environmentally dynamic marine biogeographical transition zone differs from the adjacent biogeographical provinces.

## 1 INTRODUCTION

Biogeographical transition zones are areas located between the distribution ranges of two or more independently-evolving biotic entities (e.g. biogeographical regions, provinces, biomes or communities), whose intermediate environmental conditions allow overlap of faunal elements, but prevent the extension of their ranges into each other (Ferro & Morrone, 2014). They may be found in locations with either strong environmental gradients or ‘ribbons’ of relatively unsuitable habitats (Glor & Warren, 2011), and often have high biodiversity because of the co-occurrence of species from the adjacent biogeographical entities (Morrone, 2006; Ortega & Arita, 1998; Silva-Pereira et al., 2020). On the other hand, as the gradients in environmental and associated ecological conditions represent ‘filters’ that limit the dispersal of biotic components into the transition zone and beyond (Simpson, 1965), biodiversity may be low when conditions in the transition zone are challenging for species from the neighbouring species assemblages (Pielou, 1992).

Many species exist whose ranges span multiple biogeographical units and the transition zones separating them (Teske et al., 2011). In these species, environmental clines are often mirrored by genetic clines (Riginos et al., 2011; Sotka et al., 2004; Teske et al., 2011). This suggests that such widespread species in fact comprise multiple unique, regional populations that are each on their own evolutionary trajectory, making biogeographical transition zones interesting areas for exploring incipient speciation and its environmental drivers (Dawson, 2005; Sommer et al., 2014; Teske et al., 2019).

While many studies have explored the role of biogeographical transitions in separating populations or species, the question whether the unique environmental conditions within transition zones may harbour unique biodiversity has received comparatively little attention. Some transition zones, including the area around Wallace’s Line or the South American transition zone, can be extensive and harbour endemic species (Ferro & Morrone, 2014; Mayr, 1944; Tänzler et al., 2014), but these are often difficult to study because they have complex geological or climatic histories (Darlington, 1957; Woodruff, 2003), and they may themselves contain multiple smaller-scale biogeographical units (Ferro & Morrone, 2014).

Marine biogeographical transition zones that are located on linear coastlines tend to have environmental gradients and associated biotic patterns that are particularly well defined.. Examples of such transition zones have been reported from the south-eastern Pacific (Hormazabal et al., 2004; Tapia et al., 2014), California and Florida (Pelc et al., 2009), northwestern France (Gallon et al., 2014) and along the coast of South Africa (Teske et al., 2011). These typically have well-defined gradients in sea-surface temperature, salinity and other environmental parameters.

In marine transition zones where species turnover is associated with abrupt environmental changes or unsuitable habitat, limited space makes the presence of endemic species unlikely. The present study explored the issue of endemicity in a more extensive marine biogeographical transition zone, the Wild Coast in eastern South Africa. This transition zone separates the subtropical Natal province on the east coast from thewarm-temperate Agulhas province on the south coast, and is characterised by a north-to-south temperature gradient resulting from the warm, southward-flowing Agulhas Current being gradually deflected away from the coast by the widening continental shelf. The transition zone is extensive and stretches over several hundred kilometres, although there is disagreement concerning the exact boundaries between the marine biogeographical provinces in this area (Lombard, 2004; Spalding et al., 2007; Teske et al., 2011; von der Heyden, 2009). Based on species turnover, the distinctness of this region’s fauna is only weakly supported (Emanuel et al., 1992; Turpie & Clark, 2007), and there are no prior records of endemic species (Jooste et al., 2018). This study reports the first evidence for genetically distinct populations whose ranges are limited to the Wild Coast transition zone.

## 2 METHODS

The study species, *Callichirus kraussi* (Stebbing, 1900), commonly known as sandprawn, is a widespread thalassinid decapod crustacean that ranges from the cool-temperate west coast on the Atlantic Ocean to the tropical south-western Indian Ocean (Branch et al., 2010; Teske et al., 2009). It comprises at least four phylogenetically-distinct evolutionary lineages whose ranges are confined to the region’s temperature-defined marine biogeographical provinces (Teske et al., 2009), suggesting that these could be uniquely-adapted cryptic species. Although the sandprawn’s dispersal potential is low because the adults live in burrows and the non-planktonic larvae grow up in the parent burrow (Forbes, 1973), this species is exceptionally common, and one of the dominant intertidal invertebrates in the lower reaches of the region’s estuaries and sheltered marine habitats (Day, 1981; Hanekom et al., 1988; Hanekom & Russell, 2015).

A total of 306 sandprawns were collected from 13 sites in eastern South Africa, a sampling range that includes the eastern portion of the Agulhas province, the Wild Coast transition zone and the complete Natal province (Table 1). After removal of the smaller of the two first chelae, the prawns were released. Muscle tissue from the chela was immediately placed into CTAB extraction buffer containing proteinase K, and the extraction procedure was completed in the laboratory using the CTAB protocol (Doyle, 1991). A portion of the mitochondrial cytochrome oxidase c subunit I (COI) was amplified using forward primer CrustCOIF and reverse primer PeracCOIR, as described previously (Teske et al., 2009). Sequences were aligned in MEGA7 (Kumar et al., 2016) and trimmed to a length of 570 bp.

**TABLE 1.** Estuaries on the south-east coast of South Africa (from north to south) where specimens of *Callichirus kraussi* were collected, including GPS coordinates and number of mtDNA COI sequences generated per site *(N).* Sites were assigned to three biogeographical entities.

| Entity | Site no. | Site name | GPS coordinates | *N* |
| --- | --- | --- | --- | --- |
| Natal Province | 1 | Mzingazi (Richards Bay) | 28°47'50"S, 32°05'14"E | 29 |
|  | 2 | Warner Beach (Durban) | 30°04'41"S, 30°52'19"E | 17 |
|  | 3 | Mtentweni (Port Shepstone) | 30°42'32"S, 30°28'53"E | 9 |
|  | 4 | Mpenjati (Port Edward) | 30°58'02"S, 30°16'35"E | 18 |
| Wild Coast | 5 | Mkozi (Cathedral Rock) | 31°26'48"S, 29°45'43"E | 20 |
|  | 6 | Mbotyi | 31°27'51"S, 29°44'06"E | 29 |
|  | 7 | Bulolo (Port St Johns) | 31°39'00"S, 29°30'59"E | 25 |
|  | 8 | Cintsa (East London) | 32°49'55"S, 28°06'50"E | 25 |
|  | 9 | Great Fish | 33°29'33"S, 27°07'43"E | 27 |
| Agulhas Province | 10 | Swartkops (Port Elizabeth) | 33°52'09"S, 25°37'45"E | 29 |
|  | 11 | Kabeljous (Jeffreys Bay) | 34°00'18"S, 24°55'51"E | 26 |
|  | 12 | Keurbooms (Plettenberg Bay) | 34°02'43"S, 23°22'41"E | 26 |
|  | 13 | Gourits | 34°20'15"S, 21°52'36"E | 26 |

Genealogical relationships between COI haplotypes were assessed by constructing a minimum-spanning network in popArt 1.7 (Leigh & Bryant, 2015). The program BEAST 2.6.2 (Bouckaert et al., 2014) was used to conduct a discrete phylogeographic analysis (Lemey et al., 2009) to reconstruct the location states of ancestral branches. A maximum clade credibility tree was reconstructed using 10^8^ iterations, with trees saved every 10^4^ iterations. A COI mutation rate of 1.4% per million years was specified (Knowlton & Weigt, 1998), the HKY model (Hasegawa et al., 1985) was selected based on the Bayesian Information Criterion in MEGA, and default settings were used for all other parameters. The run was repeated twice to check for consistency of results. Following assessment of convergence and effective sample size (ESS) values in Tracer 1.7 (Rambaut et al., 2018), the first 10% of trees was removed as burnin in TreeAnnotator, and a phylogenetic tree constructed using median heights was visualised in FigTree 1.4.3 (Rambaut & Drummond, 2012). The program MEGA7 was used to calculate mean and minimum Kimura 2-Parameter distances (Kimura, 1980) between regional genetic clusters identified using the haplotype network and the maximum clade credibility tree, and these were compared with mean within-lineage distances. The data were also explored for the presence of a ‘barcoding gap’ (Meyer & Paulay, 2005) by calculating a suitable distance threshold at which the number of false positives and false negatives is lowest in Spider v1.1-5 (Brown et al., 2012). As the number of sequences for this program is limited to 200, we removed the three northern most sites and the two southernmost sites (1-3 and 12-13, Table 1), but included all migrants. Following some exploratory runs, the starting threshold was set to 0 and the end threshold to 0.01, with intermediate thresholds explored at increments of 0.0001.

## 3 RESULTS

Sequence data have been submitted to the GenBank database under accession numbers MT578899 - MT579204. Four evolutionary lineages were identified, each of which occupies a distinct portion of the coast and has a single numerically dominant haplotype (Figure 1). The Agulhas and Natal provinces each have a single lineage, and two lineages were found in the Wild Coast transition zone. Interestingly, the Wild Coast lineages were not closely related to each other (Figure 2), but are derived from the subtropical and warm-temperate lineages, respectively (Figure 3). The maximum clade credibility tree depicts a sequence of evolutionary events that started with a split into a basal northern lineage (red) and a southern lineage (blue). Each of these subsequently gave rise to a Wild Coast lineage in the region closest to it, one in the north (yellow) and the other in the south (green). Subsequent southward dispersal into adjacent regions (indicated by arrows) is evident in three cases. Mean and minimum genetic distances between the four regional evolutionary lineages were low, but in all cases exceeded within-lineage distances (Table S1, Supplementary material). The most suitable genetic distance threshold to identify a barcoding gap was 0.00088 (Fig. S1, Supplementary material).

**FIGURE 1.**
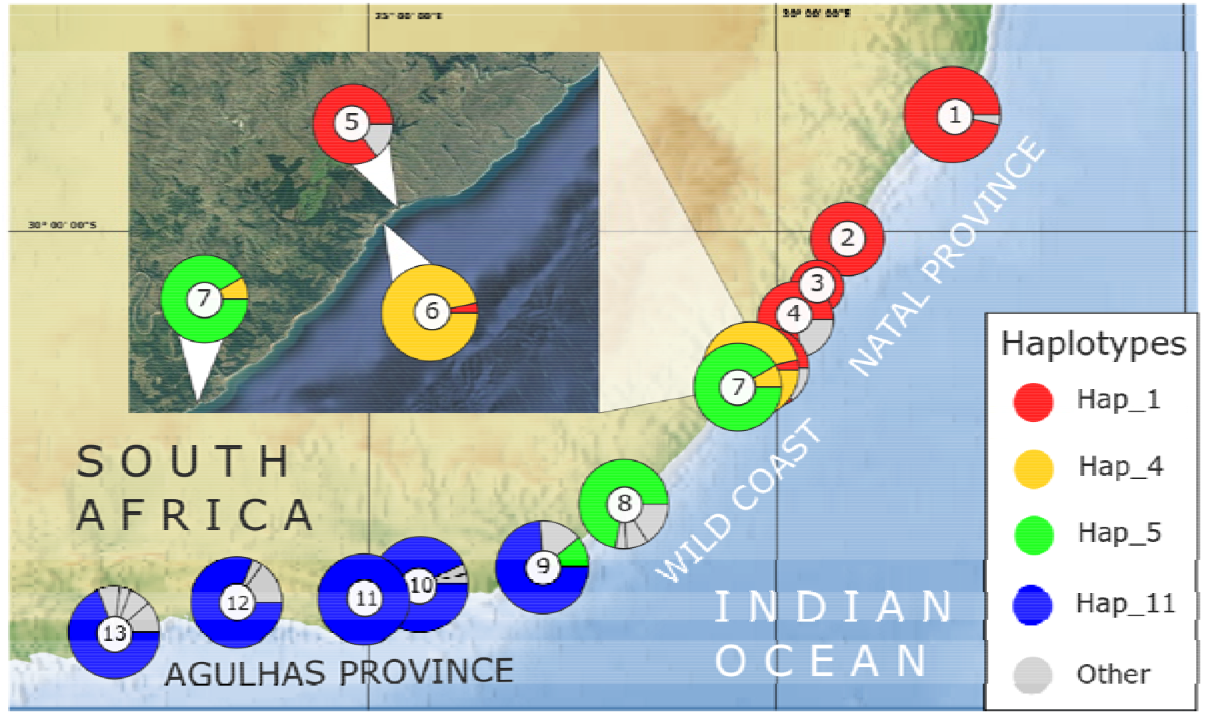
Distribution of COI haplotypes of the sandprawn *Callichirus kraussi* at 13 sites along the coastline of eastern South Africa (Table 1), with the size of each circle reflecting the sample size and the size of pie chart slices the frequency of individual haplotypes. Colours are used to distinguish the four most common haplotypes, while less frequent haplotypes are shown in grey. The insert shows a region in the northern Wild Coast where three major evolutionary lineages occur in close proximity.

**FIGURE 2.**
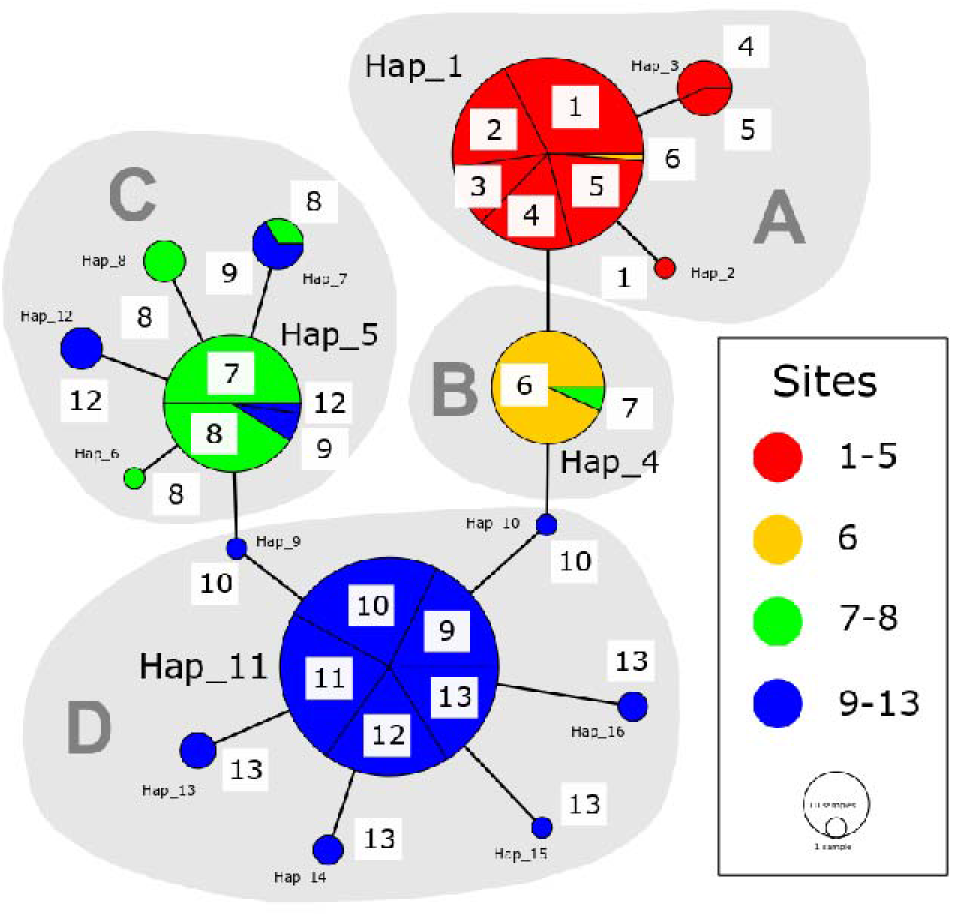
A minimum-spanning haplotype network of the COI haplotypes of *Callichirus kraussi*. Each circle represents a unique haplotype, and the size of each circle reflects the haplotype’s frequency. Connecting branches represent single nucleotide differences between haplotypes. Colours in this case were used to assign haplotypes to to four sections of coastline (Table 1). Site numbers (which correspond to those in Fig. 1 are shown in white squares. Haplotypes were assigned to four evolutionary lineages (A-D). These closely match the four sections of coastline, as each lineage is dominated by haplotypes from a specific biotic entity (A: Natal Province, B: Northern Wild Coast, C: Central Wild Coast, D: Agulhas Provinces), but the lineages also include migrants from other regions.

**FIGURE 3.**
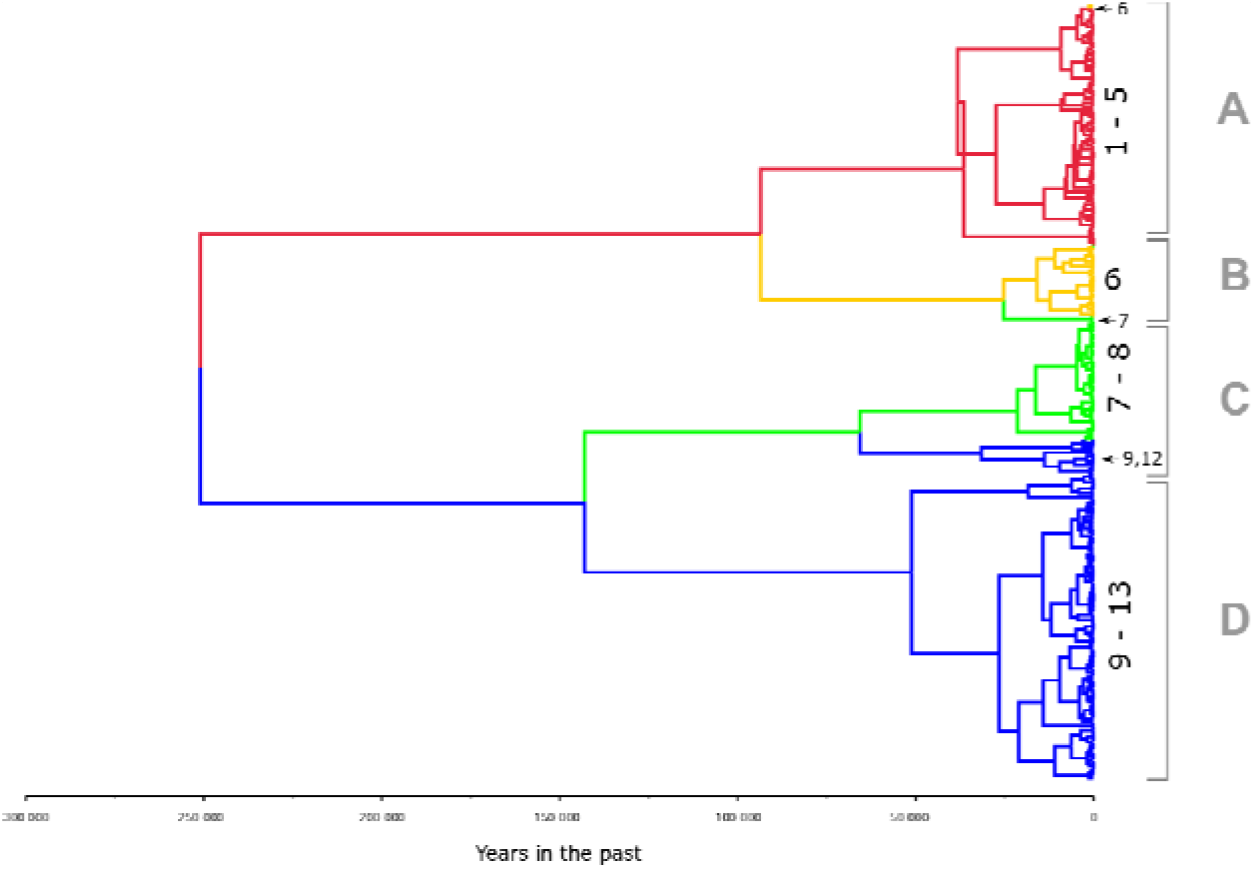
Bayesian maximum clade credibility tree constructed from COI sequences of the sandprawn *Callichirus kraussi*. Colours represent four sections of coastline (see Fig. 2), with internal branches indicating the most likely geographical location of a branch in the past, and the scale at the bottom indicating the approximate time of divergence. Four evolutionary lineages were identified, whose ranges are centred on a specific portion of the coastline (A: Natal Province, B: Northern Wild Coast, C: Central Wild Coast, D: Agulhas Province), but which also include migrants from other regions (indicated by arrows), i.e. individuals that are nested within lineages that are numerically dominated by individuals from other regions.

## 4 DISCUSSION

Marine transition zones are often located in areas where oceanographic conditions may create steep environmental gradients that are intermediate between those of the biogeographical regions they separate. For many species, they represent range edges where abiotic conditions are at the extremes of their environmental niche tolerance ranges (Sagarin & Gaines, 2002). With the majority of organisms that disperse into these regions being physiologically maladapted, there are clear selective advantages for local populations to adapt to these conditions, but the identification of endemic biodiversity is not usually a primary aim when studying marine biogeographical transitions. Comprehensive taxonomic or genetic research can potentially elevate the status of transition zones from areas of range overlap to regions whose unique biodiversity make them priority areas for management and conservation.

An example of another African marine biogeographical transition zone that is located on the south-west coast of South Africa, and which separates cool-temperate Atlantic Ocean fauna from warm-temperate Indian Ocean fauna, presents a case in point. While older literature considered it to be a region of overlap (Brown & Jarman, 1978; Day, 1969), it is now treated as a distinct coastal biogeographical region, the “South-western Cape Bioregion” (Lombard, 2004). This status is also supported by the presence of local, genetically-distinct populations of more widespread species that may represent cryptic species (Teske et al., 2011; von der Heyden, 2009).

The Wild Coast marine biogeographical transition zone has similarly been treated as a region of faunal overlap (Day, 1981; Harrison, 2002), although some studies, using community analysis, have found weakly differentiated clusters (Emanuel et al., 1992; Turpie & Clark, 2007). Similarly, most genetic studies have found extensive range overlap between sister lineages of temperate and subtropical species in this region (Teske et al., 2008; Zardi et al., 2007). Evidence for endemicity was so far limited to distinct genetic clusters rather than taxonomically-described morphospecies (Jooste et al., 2018), but, even for these, the evidence was not strong. The rocky shore limpets *Scutellastra barbara* and *S. longicosta* had south-east coast lineages whose ranges approximately matched those of the *C. kraussi* lineage from the central and southern Wild Coast (Mmonwa et al., 2015). However, the fact that the area sampled for both species excluded the subtropical Natal Province, where both are present (Branch et al., 2010), suggests that the Wild Coast may merely be the southern distribution limit of more widespread evolutionary lineages. The prawn *Palaemon capensis* shows unique mtDNA-based population structure in the southern Wild Coast, but no distinct lineages were found, and this result may be an artefact of range-wide isolation by distance (Wood et al., 2017). At the southern edge of the transition zone, a genetically-distinct site was found in the brown mussel *Perna perna* using microsatellite data, but this was not confirmed with mtDNA sequence data (Ntuli et al., 2020). Similarly, a unique Wild Coast cluster of genomic loci under thermal selection was found in an endemic goby (Teske et al., 2019), but again, this region did not have a distinct mtDNA lineage (Drost et al., 2016). Mean genetic distances between the four regional evolutionary lineages of *Callichirus kraussi* identified in this study are below commonly used fixed distance thresholds of ~0.01-0.02 for DNA barcoding of crustaceans (Bezeng & van der Bank, 2019; Raupach et al., 2015). Based on this criterion, the lineages would thus not be considered to be distinct species. On the other hand, the fact that within-lineage genetic distances were consistently smaller than minimum between-lineage distances indicates the existence of a ‘barcoding gap’ (Meyer & Paulay, 2005), and justifies the application of a lower distance threshold to delineate populations. This, and the clearly defined geographical ranges of each of the four lineages, provides the first evidence for endemic diversity within the Wild Coast transition zone.

. Some dispersal into adjacent bioregions, likely driven by the southward-flowing Agulhas Current, was evident, and exploring whether individuals from different mtDNA lineages that were found at the same sites can hybridise presents an interesting future endeavour. *Callichirus kraussi* is not a suitable candidate for traditional taxonomy because its morphology is highly conserved, and even individuals from the species’ genetically highly-divergent tropical population (Teske et al., 2009) are not distinguishable from the south coast population (Peter Dworschak, Naturhistorisches Museum Wien, pers. comm.). However, in addition to the mtDNA-based genetic differences identified here, there may be physiological adaptations that differ between populations and limit the amount of admixture between regions. For example, subtropical sandprawns can osmoregulate more efficiently than their warm-temperate counterparts, particularly at lower salinities (Cerff, 1986; Forbes, 1974), which may be an adaptation to the combination of greater precipitation and less seawater input into east coast estuaries (Teske et al., 2009). Wild Coast prawns may have intermediate osmoregulatory abilities, in addition to being adapted to a unique thermal environment, as shown for other species from this region (Papadopoulos & Teske, 2014; Teske et al., 2008, 2019; Zardi et al., 2011). Hence, although mtDNA-based divergence of the Wild Coast populations is minimal, gene regions that are involved in environmental adaptation may already be much more divergent (Teske et al., 2019).

The data generated here are not considered to be suitable for inferring what oceanographical conditions existed along the Wild Coast during the time when the regional populations split from their sister taxa. This is because mtDNA is often under strong selection (Meiklejohn et al., 2007; Stewart et al., 2008) and thus violates the assumption of the neutral theory of evolution (Kimura, 1983), which makes its usefulness for molecular dating questionable (Matumba et al., 2020). Our maximum clade credibility tree indicates that the splits may have occurred during the previous interglacial phase (~120 000 years ago) or during the subsequent glacial phase, although potential diversifying selection linked to thermal selection suggests that divergence could have taken place more recently. Genetic patterns that evolved earlier are nonetheless likely maintained by contemporary environmental conditions. The rare genetic lineage that was dominant at site 6 is likely affected by a semi-permanent cyclonic eddy reported in this area (Roberts et al., 2010), which may limit long-distance dispersal. Its location south of a coastal indent that limits the impact of the warm Agulhas Current may result in reduced influx of migrants from the northern lineage, while also reducing nearshore temperatures. Similarly, mixing between the southern Wild Coast lineage and its warm-temperate sister lineage may be limited by the widening of the continental shelf in this area, which reduces the direct influence of the Agulhas Current, and generates a nearshore countercurrent and strong upwelling (Lutjeharms et al., 2000).

### 4.1 Conclusion

Despite its importance as a biogeographical transition zone that limits the southward dispersal of Western Indian Ocean fauna into the temperate regions at the southern tip of Africa, the Wild Coast has received comparatively little scientific attention. The present study represents a significant advancement in that regard, and rejects the finding of a previous study that tentatively rejected the idea that the Wild Coast has endemic biodiversity (Jooste et al., 2018). Given the very small range of the northern Wild Coast lineage, a fine-scale sampling approach is clearly required to properly document the biodiversity of this poorly-studied region, and determine whether the spatial genetic patterns found here are unique to *C. kraussi* or represent a more general phenomenon. Several estuarine species of high conservation priority that occur elsewhere in South Africa have very small distribution ranges (Allanson, 1958; Penrith & Penrith, 1972; Whitfield, 1995; Whitfield et al., 2017), and it is likely that such species also exist along the Wild Coast.

